# *In Vivo* Proximity Labeling Identifies a New Function for the Lifespan and Autophagy-regulating Kinase Pef1, an Ortholog of Human Cdk5

**DOI:** 10.1101/2024.06.12.598664

**Authors:** Haitao Zhang, Dongmei Zhang, Ling Li, Belinda Willard, Kurt W. Runge

**Author notes:** Correspondence to Kurt W. Runge.

## Abstract

Cdk5 is a highly-conserved, noncanonical cell division kinase important to the terminal differentiation of mammalian cells in multiple organ systems. We previously identified Pef1, the *Schizosaccharomyces pombe* ortholog of cdk5, as regulator of chronological lifespan. To reveal the processes impacted by Pef1, we developed APEX2-biotin phenol-mediated proximity labeling in *S. pombe.* Efficient labeling required a short period of cell wall digestion and eliminating glucose and nitrogen sources from the medium. We identified 255 high-confidence Pef1 neighbors in growing cells and a novel Pef1-interacting partner, the DNA damage response protein Rad24. The Pef1-Rad24 interaction was validated by reciprocal proximity labeling and co-immunoprecipitation. Eliminating Pef1 partially rescued the DNA damage sensitivity of cells lacking Rad24. To monitor how Pef1 neighbors change under different conditions, cells induced for autophagy were labeled and 177 high-confidence Pef1 neighbors were identified. Gene ontology (GO) analysis of the Pef1 neighbors identified proteins participating in processes required for autophagosome expansion including regulation of actin dynamics and vesicle-mediated transport. Some of these proteins were identified in both exponentially growing and autophagic cells. Pef1-APEX2 proximity labeling therefore identified a new Pef1 function in modulating the DNA damage response and candidate processes that Pef1 and other cdk5 orthologs may regulate.

## INTRODUCTION

The fission yeast cyclin-dependent kinase Pef1 belongs to the conserved cdk5 kinase family, with conservation so strong across evolution that human cdk5 can complement its fungal counterparts even while interacting with fungal cyclins (1,2). In humans, cdk5 dysregulation is associated with neurodegenerative disorders such as Alzheimer and Parkinson disease, as well as various solid and hematological cancers (3). In a previous study, we found that Pef1-Clg1 Cdk-Cyclin kinase complex negatively regulates chronological lifespan, a process that measures the ability of cells to remain viable during starvation-induced quiescence (4). cdk5 has also been implicated in modulating autophagy in both mammalian cells and *Drosophila melanogaster* by phosphorylating different substrates (5–7). Genetic experiments in *Schizosaccharomyces pombe* ((8) and our unpublished data) showed that fission yeast Pef1 exerts a negative role in macroautophagy, a process induced by starvation that recycles cellular proteins and components (9). Increased levels of autophagy and extended lifespan during starvation in cells lacking Pef1 are consistent as the nutrients provided by autophagy can extend the viability of starved cells. However, the underlying mechanism(s) by which Pef1 regulates autophagy and lifespan remains unclear. Identifying the upstream and downstream molecules that affect Pef1 activation and signaling would elucidate how Pef1 regulates cell physiology. Given that autophagy is an inducible process, an inducible system that can identify the upstream and downstream proteins near Pef1 under different conditions would be extremely useful.

Traditional affinity purification coupled with mass spectrometry analysis (AP-MS) (10) enables the identification of protein partners of interest, providing catalogs of the functions and biological processes associated with the bait protein. However, affinity purification primarily captures stable physical interactions and potentially misses transient binding between kinases and their substrates, as well as interactions disrupted during sample preparation. The development of proximity labeling (PL) in living cells addresses these shortcomings. PL utilizes an engineered enzyme fused to a bait protein to covalently tag nearby proteins within nanometer distances (11,12).

One promising PL approach involves fusing the bait protein with a promiscuous biotin ligase, such as BioID and BioID2 (13,14). These enzymes convert biotin to biotin-AMP which reacts with the unmodified lysines of nearby proteins. However, the slow enzyme kinetics of BioID and BioID2 requires at least 18 hours to get sufficient labeling, making them unsuitable for studying kinase interactomes and transient interactions. Moreover, their optimal activity at 37°C renders them inactive in yeast. A biotin ligase engineered through yeast display-based directed evolution that works well at temperature as low as 30°C, TurboID and miniTurboID, exhibits significantly increased enzyme activity, achieving equivalent biotinylation in 10 minutes compared to the 18-hour requirement of BioID/BioID2 (15). To some extent, the labeling by TurboID and miniTurboID can be considered inducible as the K_M_ for biotin is higher than physiological biotin concentrations, and biotinylation can be induced by treating cells with high levels of biotin in the medium (15). However, it is important to note that the constitutive enzyme activity of biotin ligase can result in background biotinylation (15,16), potentially due to different concentrations of biotin in different subcellular locations.

The other candidate for proximity labeling is the engineered soybean ascorbate peroxidase APEX2, known for its efficacy across a wide temperature range and its activity in fixed samples (11,17–19). Upon activation by a 1-minute treatment with 1 mM H_2_O_2_, APEX2 rapidly converts the biotin-phenol (BP) substrate to a short-lived (<1 ms) biotin-phenoxy radical, which covalently conjugates to tyrosine, lysine and cysteine residues of proteins within 20 nm (20). APEX2 remains inactive in the absence of H_2_O_2_ and exhibits negligible background activity. Its inducible and rapid labeling reaction makes it suitable for capturing interactions occurring under special conditions. Consequently, APEX2 has found widespread use in mapping the protein-protein and protein-DNA/ RNA networks associated with DNA damage repair (21), a predefined genomic locus (22,23), ER-to-Golgi transport (24) and G-protein-coupled receptors (GPCRs) (25). While concerns have been raised about the effect of the 1 mM H_2_O_2_ pulse on protein localization (15), we note that the 1 mM H_2_O_2_ pulse is short and at a level in between H_2_O_2_ concentrations used in longer treatments to induce mild or acute oxidative stress (0.3 to 3 mM for ≥ 5 minutes) (26–28). The combination of relatively low H_2_O_2_ concentration, the short exposure and past success of this approach indicated that APEX2-labeling would be a fruitful approach to monitor rapidly inducible processes in *S. pombe*.

The application of PL to map yeast proteomic networks has been limited. The working temperatures of BioID/BioID2 renders this approach impractical in yeast cells. TurboID has been recently applied to fission yeast; however, the biotin in both rich and minimal media is sufficient for TurboID to biotinylate adjacent proteins (16). Without biotin starvation, TurboID did not respond to addition of biotin. TurboID requires at least 30 minutes to get sufficient labeling in cells pre-starved with biotin, thereby diminishing its “inducible” attribute observed in mammalian cells and potentially altering the cell physiology as well.

A challenge in using APEX2 in yeast cells arises from the inability of BP to permeate the cell wall. A previous study (29) reported that treatment of yeast cells with Zymolyase-100T or high osmotic solutions (1.2 M sorbitol and 1 M KCl) initiated APEX2-dependent biotinylation from a fusion protein located near the plasma membrane, suggesting that APEX2-based proximity labeling can be used in fission yeast. Our study tested this technique by mapping the neighborhood of Pef1 in both exponential and autophagic cells. While these previously reported treatments did not allow labeling by Pef1-APEX2, we established conditions that allowed efficient biotinylation of nuclear and cytoplasmic proteins and tested whether immunodepleting the endogenous biotinylated proteins prior to mass spectral analysis changed the number of proteins detected by Pef1-APEX2 biotinylation. We identified the 14-3-3 protein Rad24 as a novel interacting partner of Pef1, and found that eliminating Pef1 partially rescued the sensitivity of cells lacking Rad24 to DNA damaging agents. Pef1-APEX2 successfully labeled cells induced for autophagy and identified proteins in actin cytoskeletal dynamics, vesicle-mediated transport and the yeast lysosome (vacuole). These proximity labeling methods for Pef1 thus identified a new Pef1 role in the response to DNA damage and new pathways that Pef1 and cdk5 orthologs in other organisms may regulate.

## MATERIALS AND METHODS

### Growth media and yeast strains

All *S. pombe* stains used in this study are summarized in Supplemental Table 1. Cells were cultured in either YES or EMMG media supplemented with appropriate nutrients to complement cellular auxotrophies unless indicated otherwise (30). Information regarding gene structures, mutant strain phenotypes and interacting proteins were accessed from Pombase (31,32) and Ensembl Fungi (33).

### Epitope tagging of endogenous biotinylated proteins

The endogenously biotinylated proteins in *S. pombe* and other eukaryotes are pyruvate carboxylase (encoded by *S. pombe pyr1^+^*) and acetyl CoA carboxylase (encoded by *cut6^+^*). To tag these proteins with 13 copies of the Myc epitope, a DNA fragment containing approximately 500 bp of genomic sequence immediately upstream of the stop codon, followed by the 13Myc epitope sequence with a stop codon, and then approximately 500 bp of genomic sequence immediately downstream of the stop codon including the transcription terminator of the gene, was assembled into the Kpn I/Xba I site (for *cut6*) or Kpn I/Not I site (for *pyr1*) of the yeast integrating plasmid pJK210 (34). The plasmids pJK210-pyr1-13Myc-*ura4^+^* and pJK210-cut6-13Myc-*ura4^+^* were verified by DNA sequencing, linearized using Nco I (which cuts in homology to *pyr1^+^* or *cut6^+^*), and transformed into KRP1 (*h^-^*) and KRP2 (*h^+^*) cells, respectively. Positive *ura4^+^* markers were confirmed by colony PCR and Western blotting before being mated with each other on SPA plates. The resulting tetrads were digested with glusulase and analyzed by random spore analysis. Strains expressing both *cut6^+^-13Myc* and *pyr1^+^-13Myc* as the only copies of these genes were identified by colony PCR and confirmed by Western blotting.

### APEX2-3V5 and 13Myc fusions to Pef1, Rad24 and Cek1

Pef1, Rad24 and Cek1 proteins were C-terminally tagged with either APEX2-3V5, 3V5 or 13Myc at the C-terminus using pFA6a-based vectors (35,36), the APEX2-3V5 and 3V5 vectors were made by ourselves and will be distributed through the Japanese National BioResource Project (NBRP https://yeast.nig.ac.jp/yeast/top.xhtml). For each clone, a plasmid comprising approximately 300 bp of genomic sequence directly upstream of the stop codon, the epitope tag fused to the open reading frame and ending with a stop codon and transcription terminator, a selectable marker and approximately 300 bp of genomic sequence downstream of the stop codon was constructed and verified by DNA sequencing. This DNA fragment was transformed into wild type cells, selecting for the marker linked to the epitope tag. The positive colonies were confirmed by colony PCR and Western blotting. The *rad24Δ* strains were generated by transforming a dominant selectable marker flanked by 300 bp of genomic sequence directly upstream of the start codon and 300 downstream of the stop codon. The *rad24Δ* deletion was validated with colony-PCR.

### In vivo proximity labeling with APEX2 and biotin-phenol (BP)

The labeling procedure was adapted from the protocol originally developed for mammalian cells (11). Cells were grown in 50 mL EMMG medium supplemented with appropriate nutrients overnight to mid log phase (approximate cell density at 4-5 x 10^6^ cells /mL). Cells were spun down, washed with 20 mL MilliQ water, and 2 x 10^8^ cells were aliquoted into each 1.5 mL labeling tube, pelleted and suspended in 500 µL of 1.2 M sorbitol and 10 mM PIPES (pH 7.0) supplemented with Zymolyase-100T (MP Biomedicals, cat. # 32093) and BP (APExBIO, cat. # A8011, made as a 0.5 M stock in DMSO and stored for several weeks at -80°C) at the indicated concentrations. The suspensions were incubated at room temperature for 30 minutes on a tube rotator at ∼30 rpm (Stuart SB3) unless noted otherwise. The labeling reaction was initiated by addition of 1 mM H_2_O_2_, followed by incubation at room temperature for 1 minute. Cells were then spun down for 30 seconds at 1,000 x g. Cell pellets were immediately washed three times with 500 µL of quencher solution containing 1.2 M sorbitol, 10 mM PIPES pH 7.0, 5 mM Trolox (Sigma-Aldrich, cat. # 238813), 10 mM sodium ascorbate (Spectrum Chemical, cat. # S1349), 10 mM sodium azide and 1 mM PMSF. The cell pellets were either frozen at -80°C immediately or directly subjected to protein extraction.

### Protein extraction and Mass-Spectrometry (MS) analysis of biotinylated proteins

The Zymolyase 100-T treated cell pellets were lysed in an ice-cold basic solution containing 1.85 M NaOH and 7.4% (v/v) β-mercaptoethanol. Total proteins were precipitated by adding trichloroacetic acid (TCA) to the lysate to a final concentration of 25% (w/v), incubating on ice for 10 minutes and then spinning at 16,100xg for 10 minutes in a microfuge. The resulting pellets were washed with ice-cold 90% acetone, gently air-dried, suspended in 240 µL HEPES lysis buffer (consisting of 50 mM HEPES pH 7.5, 150 mM NaCl, 1 mM EDTA, 5 mM NaF, 1 mM Na_3_VO_4_, 1 mM PMSF, 1X protease inhibitor (Roche, catalog # 11836170001) and 5% glycerol). The cell pellets were fully disrupted by sonication with sample tubes suspended in an ice water bath (Diagenode BioruptorXL water bath, 10 seconds on and 59 seconds off for 10 cycles with maximum power output). The milky suspensions were then solubilized by adding SDS and Tris base to final concentrations of 0.5% (w/v) and 50 mM, respectively. After centrifugation at 16,100 x g for 10 minutes, the resulting supernatants were stored at -80°C for future use. Protein concentration was determined using the BCA method (Pierce, cat. # 23227).

To enrich the biotinylated proteins from a standard labeling, 800 µg of total proteins were diluted with HEPES lysis buffer to a final volume of 1.2 mL and then incubated with 80 µL of 20% magnetic streptavidin beads (Promega, cat. # V7820) overnight at 4°C on a tube rotator.

For samples to be immunodepleted of Pyr1-13Myc and Cut6-13Myc, 1.6 mg of total proteins were diluted with HEPES lysis buffer to a final volume of 2.4 mL and incubated overnight with 80 µL of 50% anti-c-Myc agarose beads (Sigma-Aldrich, A7470) on a tube rotator. The following morning, the anti-c-Myc beads were pelleted by spinning at 1000 x g for 2 minutes, and the supernatant was transferred to a new tube and incubated with 80 µL of 20% magnetic streptavidin beads overnight as above.

The proteins on streptavidin beads were sequentially washed twice with 1 mL of RIPA buffer (50 mM Tris•HCl pH 7.5, 150 mM NaCl, 0.1% SDS, 0.5% sodium deoxycholate, 1% Triton X-100), once with 1 M KCl, once with 0.1 M Na_2_CO_3_, once with 1.2 M urea and once more with RIPA buffer. A final wash with 1 mL 50 mM HEPES pH 7.4 was performed to remove chemicals that could interfere with trypsin digestion. On-bead trypsin digestion was performed in 100 mM NH_4_HCO_3_ at 37°C overnight. The resulting peptides were desalted with C18 cartridges and dried with a Speedvac vacuum concentrator.

For LC-MS/MS analysis, the peptides solubilized in 1% acetic acid were analyzed on Thermo Scientific Orbitrap Fusion Lumos Tribrid Mass Spectrometer. Peptides were loaded onto a C18 reversed-phase columns (Dionex 15 cm length, 75 µM inner diameter, 2 µM, 100 Å) and eluted by an acetonitrile/0.1% formic acid gradient at a flow rate of 0.3 µL per minute. The eluates were introduced into the source of the mass spectrometers on-line. The micro-electrospray ion source is operated at 2.5 kV. The digest was analyzed using the data dependent multitask capability of the instrument. The full scan mass spectra were to determine peptide molecular weight and product ion spectra were to determine amino acid sequence. The data were analyzed by using all CID spectra to search the *Schizosaccharomyces pombe* SwissProt database with software Sequest.

For TMT-mass spectrometry analysis, the desalted and dried peptide digests were reconstituted in 50 mM TEAB (triethylammonium bicarbonate) solution and quantitated with Quantitative Colorimetric Peptide Assay kit (Pierce, cat. # 23275). The peptide samples from different experiments were labeled with TMT10plex tags. The labeled samples were then mixed together with the same percentage of the total peptide amount of each sample. The combined mixture of TMT-labeled tryptic peptides was desalted, dried and reconstituted in 30 µL of 1% acetic acid for LC-MS3 analysis. Five microliters of the mixture were fractionated by a reverse-phased capillary chromatography column (Dionex 15 cm length x 75 µM inner diameter Acclaim Pepmap C18, 2 µM, 100 Å). Fractionation of peptides were carried out by an acetonitrile/0.1% formic acid gradient at a flow rate of 0.3 µL per minute. The flow-through were introduced into the source of the mass spectrometers in-line. The micro-electrospray ion source was operated at 2.5 kV. The full scan mass spectra and MS2-CID spectra successively acquired by the data dependent multitasking capability of the instrument was used to determine peptide molecular weights and amino acid sequence, respectively. The MS3-HCD spectra acquired by the Synchronous Precursor Selection (SPS) capability of the instrument was used to quantitate the reporter ions. All CID spectra were searched against the *S. pombe* UniprotKB protein sequence database. False discovery rate of protein was set at 1%.

### Immunoprecipitation

To test co-immunoprecipitation between Pef1 and Rad24, cells were mechanically broken with glass beads on Precellys tissue homogenizer (Bertin Instruments) in HEPES lysis buffer supplemented with 1% (v/v) Triton-X100. A total protein of 800 µg was incubated overnight with 40 µL of anti-V5 agarose beads (Sigma-Aldrich, cat. #A7345). After incubation, the beads were washed with 1 mL of HEPES lysis buffer for 5 times. The eluates were analyzed by Western blotting with antibodies to the epitope tags (described above).

### Western blotting

The protein samples were prepared with the same method to that used for proteomic analysis described above. A total of 40 µg protein of each sample was mixed with SDS-loading buffer, separated on 4–20% precast polyacrylamide gel (Bio-rad, cat. # 4561096), transferred onto nitrocellulose membrane (Li-Cor, cat. # 926-31090) following the instructions for the Bio-rad precast gel system and probed with the indicated antibodies as previously described (37). Gels were run with prestained markers (Li-Cor, cat # 928-40000) and after transfer, the membranes were cut to separate the region with the loading control proteins and the proteins of interest, and each membrane piece was processed with the appropriate detection reagents. The primary and secondary antibodies used in this project were rabbit anti-V5 (Proteintech, cat. #14440-1-AP), mouse anti-c-Myc (Sigma-Aldrich, cat. # M4439), rabbit anti-GFP (Abcam, ab6556), rabbit anti-Histone H3 (abcam, ab1791), IRDye 800CW donkey anti-rabbit IgG (Li-Cor, cat. # 926-32213) and IRDye 680 Donkey anti-mouse IgG (Li-Cor, cat. #925-68072). Biotinylated proteins were detected using IRDye 800CW Streptavidin (Li-Cor, cat. #925-32230). The blots were scanned using the Odyssey CLx infrared imaging system.

## RESULTS

### Optimization of APEX2-dependent proximity labeling in *S. pombe* cells

*S. pombe* cells expressing Pef1 with a C-terminal APEX2-3V5 fusion from the *pef1^+^*endogenous promoter at the *pef1^+^* genomic locus (Fig. 1A) were used to optimize proximity labeling. *S. pombe* cells possess a rigid cell wall that hinders BP from being incorporated into the cell. Zymolyase-100T treatment, commonly used to digest the cell wall, facilitated BP permeation through the fission yeast cell membrane (29). To assess the *in vivo* biotinylation activity of the Pef1-APEX2 fusion protein, *pef1^+^-APEX2-3V5* cells were treated with Zymolyase-100T as well as 2.5 mM BP for 1 hour. The western blotting results revealed that biotinylation signals were detected in cells treated with both 2.5 mM BP and 1 mM H_2_O_2_, but not in cells treated with either BP or H_2_O_2_ alone (Fig. 1B). As anticipated, two endogenously biotinylated carboxylases, Pyr1 (pyruvate carboxylase, 131 kDa) and Cut6 (acetyl-CoA carboxylase, 257 kDa), were detected in both control and experimental samples. These findings show that Pef1-APEX2-3V5 can biotinylate intracellular proteins in an inducible manner. Previous work indicated that a high osmotic solution, either 1.2 M sorbitol or 1 M KCl alone could facilitate BP permeation through the cell wall of *S. pombe* and initiate biotinylation by APEX2 (29). However, these two approaches did not allow protein biotinylation in Pef1-APEX2-3V5 cells (Fig. 1B, the two rightmost lanes).

**Fig. 1.**
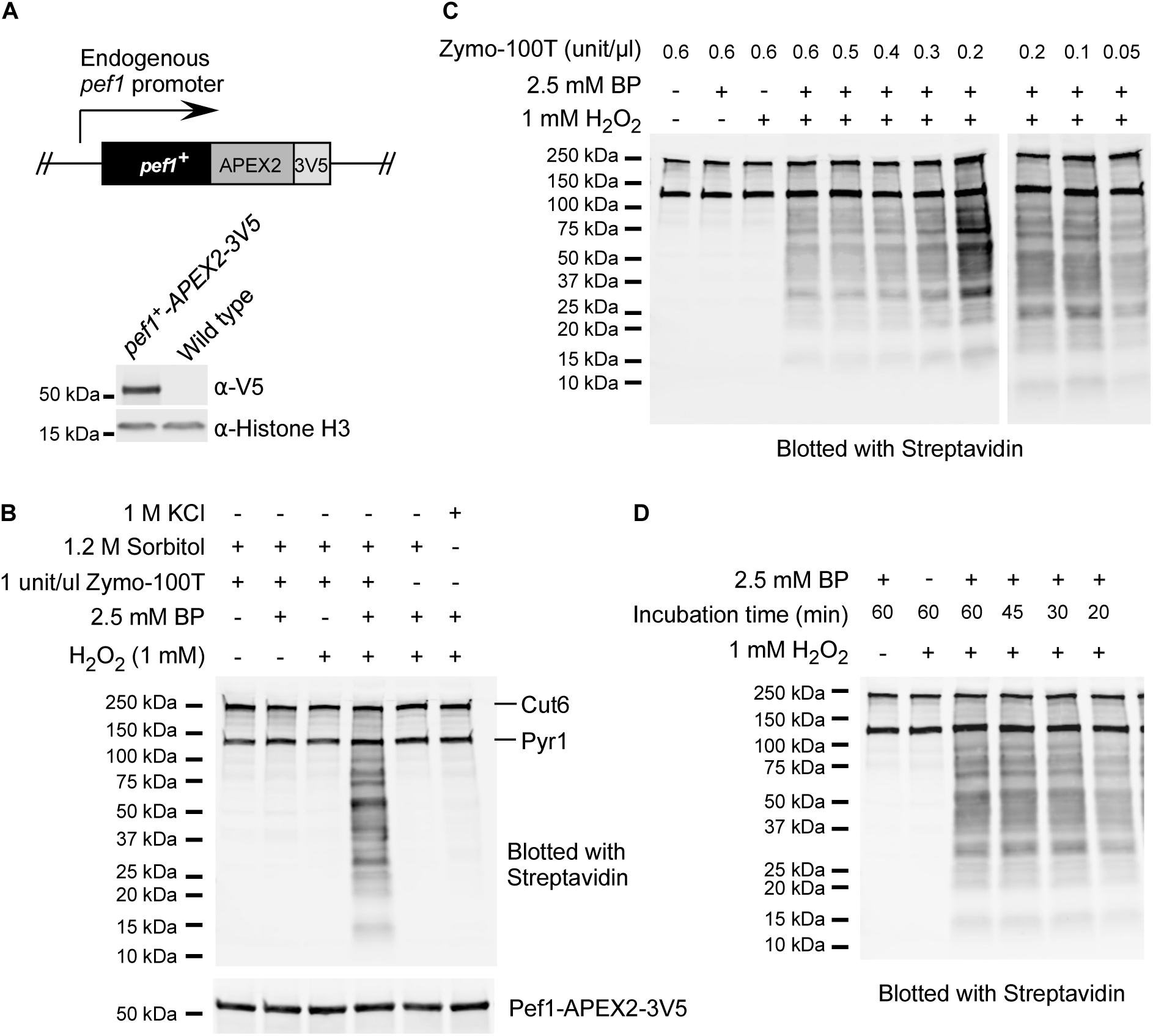
Optimization of the proximity labeling by Pef1-APEX2-3V5 in fission yeast. *(**A**)* Schematic illustration of genomic *pef1^+^* tagged with APEX2-3V5. The expression of *pef1^+^*- APEX2-3V5 under the endogenous *pef1^+^* promoter was confirmed by Western blotting using anti-V5 antibody. Histone H3 served as a loading control. *(**B**)* Pef1*-*APEX2-dependent biotinylation was detected in cells treated with Zymolyase-100T, and not in those treated with osmotic shock, by fluorescein-conjugated streptavidin. The incubation was conducted at room temperature for 1 hour before pulsed by 1 mM H_2_O_2_ for 1 minute. The endogenously biotinylated Cut6 (257 kDa) and Pyr1 (131 kDa) proteins are indicated. *(**C**)* Optimization of the concentration of Zymolyase-100T. The labeling reactions were conducted in 10 mM PIPES (pH7.0), 1.2 M sorbitol supplemented with 2.5 mM BP and different concentrations of Zymolyase-100T at room temperature for 1 hour before being treated with 1 mM H_2_O_2_ for 1 minute. The rightmost 0.2, 0.1 and 0.05 unit/µL Zymolyase-100T samples are from a separate gel. *(**D**)* Optimization of the incubation time of Zymoylase-100T. Exponential cells were incubated in 10 mM PIPES (pH 7.0), 1.2 M sorbitol supplemented with 0.2 units/µL Zymolyase-100T and 2.5 mM BP for different times at room temperature before being pulsed with 1 mM H_2_O_2_ for 1 minute. BP, H_2_O_2_ or both were omitted in negative control samples.

The incubation time and concentration of Zymolyase-100T treatment that allowed the highest labeling efficiency with the mildest treatments was then determined. The impact of different concentrations of Zymolyase-100T, ranging from 0.1 unit/µL to 0.6 unit/µL with 2.5 mM BP and a 1 hr incubation at room temperature showed that the strongest labeling signals were observed in cells treated with Zymolyase-100T at 0.2 unit/µL (Fig. 1C). Increasing the concentration of Zymolyase-100T compromised labeling efficiency, whereas 0.05 units/µL of Zymolyase-100T was insufficient for maximum labeling (Fig. 1C). While 0.1 and 0.2 units/µL gave qualitatively similar levels of labeling, the reduction in labeling at 0.05 units/µL raised concerns that 0.1 units/µL might be borderline under some growth conditions that strengthen the cell wall, and 0.2 units/µL was chosen going forward. The incubation time was optimized by comparing the intensity of biotinylation signals in cells treated with 0.2 unit/µL Zymolyase-100T and 2.5 mM BP for durations of 20, 30, 45 and 60 minutes, and showed that the biotinylation signals peaked at the 30-minute time point, and were not improved by longer incubation items (Fig. 1D). Therefore, APEX2-mediated proximity labeling was performed with 0.2 units/µL Zymolyase-100T for 30 minutes in the presence of 2.5 mM BP.

The Zymolyase-100T treatment described above was conducted in a solution of 1.2 M sorbitol and 10 mM PIPES, pH7.0, which deprived cells of both energy and nitrogen supply. We wondered whether combining EMM culture medium with Zymolyase-100T could improve maximum labeling. We tested various EMM formulations, including EMM with no nitrogen or glucose, EMM with 2% glucose, EMM plus glutamate (EMMG) and EMM with ammonium chloride (38). Each type of medium was supplemented with 1.2 M sorbitol to maintain osmolarity, and the concentration of Zymolyase-100T during labeling was varied. The concentration of BP was kept at 2.5 mM, and the incubation time was set to 1 hour at room temperature. As depicted in Fig. 2A, the biotinylation signals in cells treated with EMM medium with no nitrogen or glucose treated with 0.4 unit/µL Zymolyase-100T were comparable to those in cells treated with 0.2 unit/µL Zymolyase-100T, 1.2 M sorbitol, 10 mM PIPES, pH7.0. The addition of nitrogen source, either glutamate (Fig. 2C) or ammonium chloride (Fig. 2E), drastically decreased labeling efficiency, and increasing the amount of Zymolyase-100T did not rescue the labeling signals. Moreover, the addition of 2% glucose completely inhibited the labeling reaction (Fig. 2B and 2D).

**Fig. 2.**
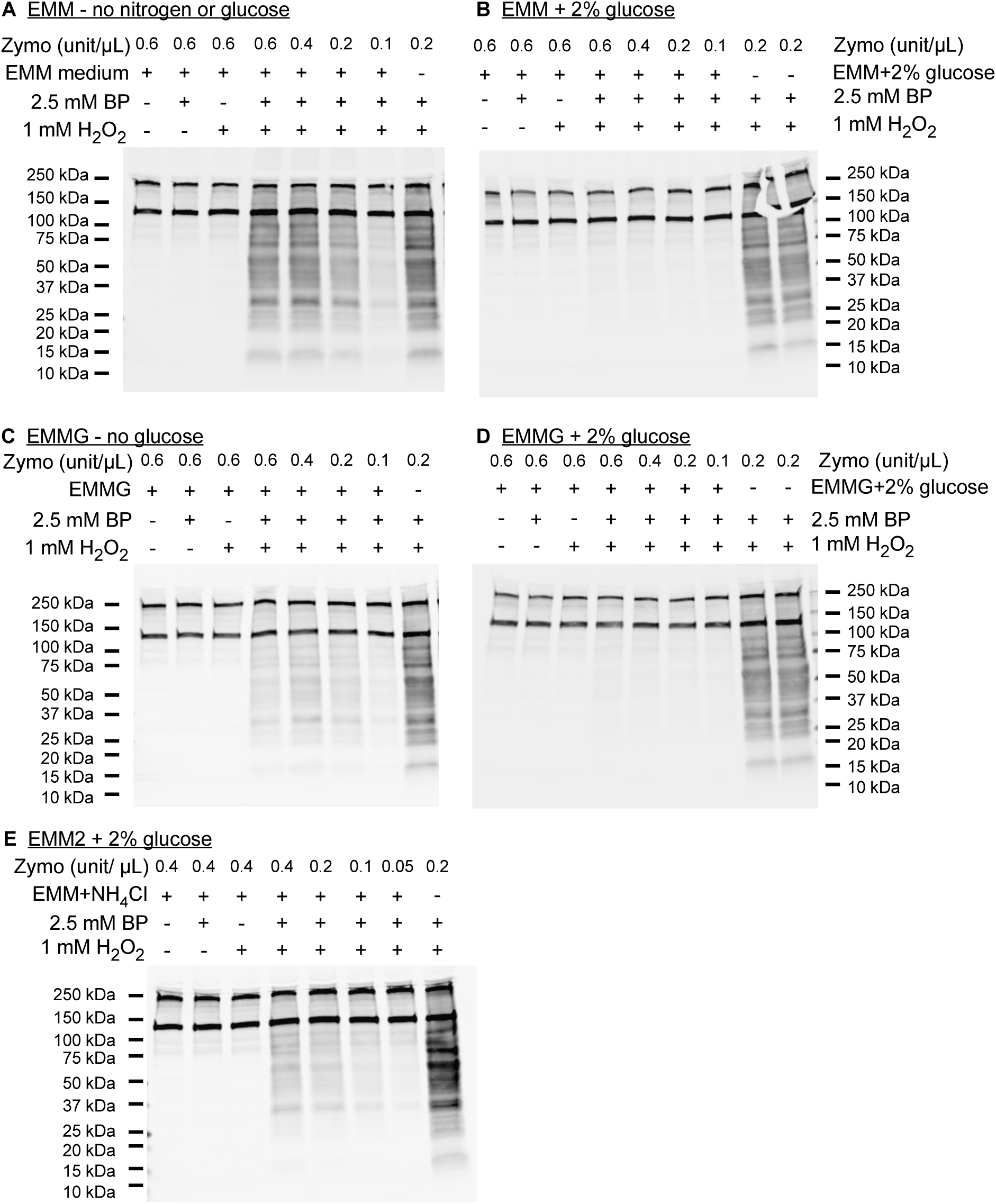
The proximity labeling mediated by Pef1-APEX2-3V5 was inhibited in the presence of a nitrogen source and 2% glucose. The proximity labeling reactions were conducted in different culture media containing 2.5 mM BP and varied concentration of Zymolyase-100T for 1 hour at room temperature, followed by a 1-minute pulse of 1 mM H_2_O_2_. *(**A**)* EMM, *(**B**)* EMM plus 2% Glucose, *(**C**)* EMMG (21.3 mM glutamate), *(**D**)* EMMG (21.3 mM glutamate) plus 2% glucose, and *(**E**)* EMM2 (93.5 mM ammonium chloride). All reaction media contained 1.2 M sorbitol to maintain cell membrane integrity. Either BP, or H_2_O_2,_ or both were omitted in control samples. Positive control samples are indicated by a “–” sign in the row showing the medium and indicate labeling in 1.2 M sorbitol plus Zymolyase and BP (see Materials and Methods). Biotinylation signals were detected with fluorescein-conjugated streptavidin.

These results indicate that the most effective proximity labeling mediated by Pef1-APEX2-3V5 occurs when cells are treated with 0.2 unit/µL Zymolyase-100T in a solution of 1.2 M sorbitol and 10 mM PIPES, pH 7.0 for a duration of 30 minutes, under the presence of 2.5 mM BP. This optimized protocol was then utilized in subsequent studies.

### Immunodepletion of endogenous biotinylated proteins Cut6 and Pyr1 did not increase the identification of Pef1 neighbors

Acetyl CoA carboxylase and Pyruvate carboxylase, Cut 6 and Pyr1 respectively, are endogenously biotinylated proteins that comprised a significant portion of the streptavidin-purified proteins and co-purified with APEX2-labeled proteins (Figs. 1, 2). As the peptides generated from these proteins might interfere with the detection of other low abundance proteins, we asked whether depleting Cut 6 and Pyr1 would allow the identification of a larger number of Pef1 neighbors. Both *cut6^+^* and *pyr1^+^* are required for cell growth. We therefore attempted to immunodeplete these protein prior to streptavidin purification. The *cut6^+^*and *pyr1^+^* coding sequences were each fused with sequences encoding 13 copies of the myc epitope (13Myc) in *pef1^+^-APEX2-3V5* cells. Notably, the 13Myc tagging still allowed cells to grow well in standard media (data not shown).

Cells expressing Pef1-APEX2-3V5 with or without the 13Myc tagged Cut6 and Pyr1 were labeled in duplicate to provide sufficient protein for immunodepletion and streptavidin purification. The presence of Cut6-13Myc and Pyr1-13Myc did not cause noticeable changes in biotinylation compared to cells lacking these tags (Fig. 3A). Both the BP-labeled and non-labeled extracts underwent immunodepletion by incubation with anti-Myc agarose beads, the supernatants were transferred to new tubes and incubated with streptavidin beads to purify the biotinylated proteins. As a control, BP-labeled and non-labeled pef1-APEX2-3V5 cells were also directly incubated with streptavidin beads without depletion. Immunodepletion removed the majority Cut6-13Myc and Pyr1-13Myc proteins as judged by Coomassie Blue staining (Fig. 3B). The streptavidin bound proteins were then subjected to on-bead digestion by trypsin. Peptides from different samples were labeled with different TMT-isobaric tags, pooled and analyzed by mass spectrometry. When compared to the respective negative controls lacking the H_2_O_2_ pulse, a total of 187 proteins were identified with at least 2 unique peptides and a MS signal of ≥2 in the non-depleted samples and 183 proteins matching these criteria were identified in the immunodepleted cells (Supplemental Table 2).

**Fig. 3.**
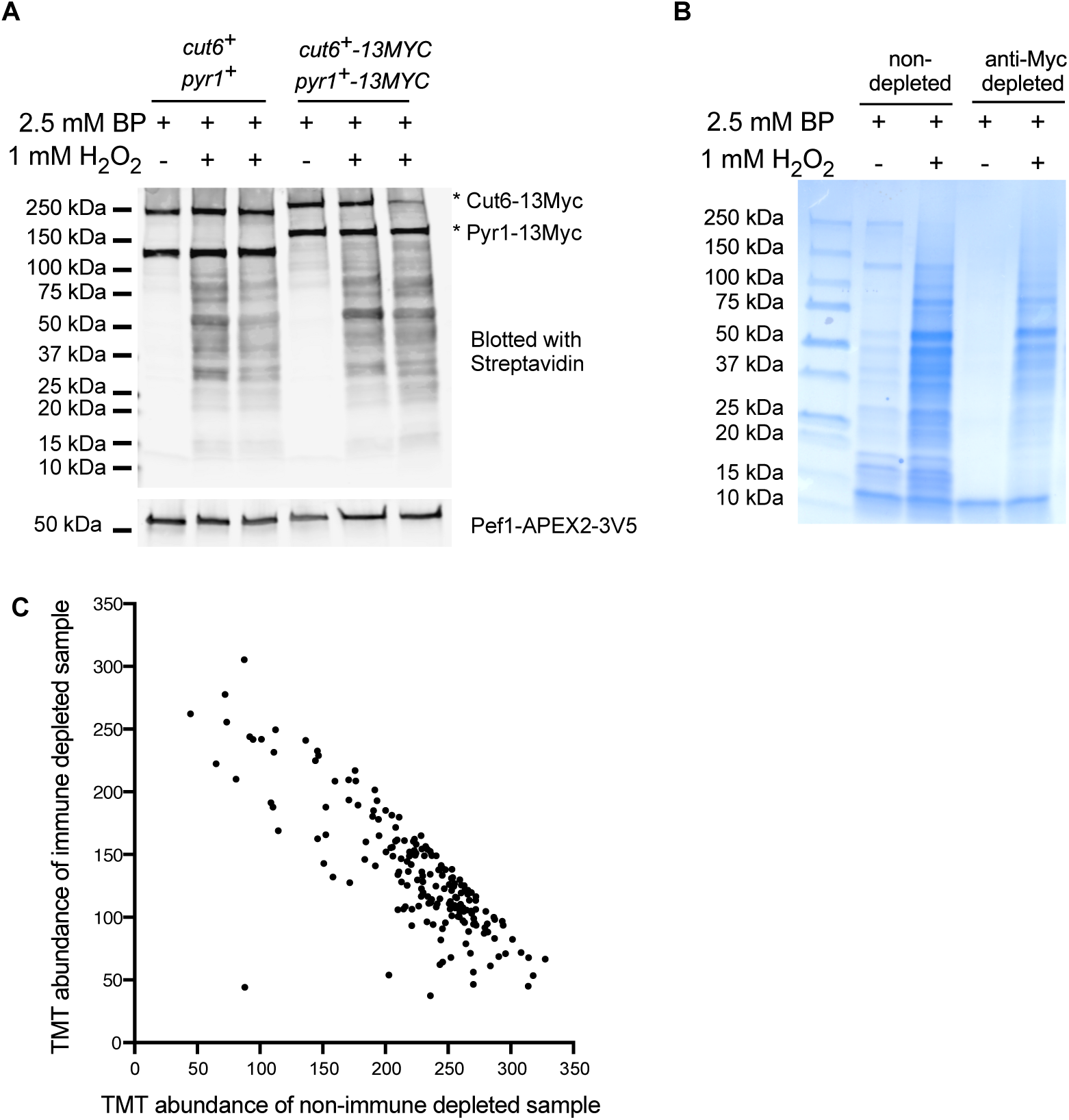
Immuno-depletion of endogenous biotinylated carboxylases did not enhance the identification of Pef1-APEX2-3V5 labeled proteins. *(**A**)* The 13Myc tagging of genomic *cut6*^+^ and *pyr1^+^*did not alter the labeling efficiency of Pef1-APEX2-3V5. Exponential *pef1*^+^*-APEX2-3V5* cells with or without 13Myc-tagged *cut6*^+^ and *pyr1*^+^ were incubated with 0.2 units/µL Zymolyase-100T and 2.5 mM BP for 30 minutes at room temperature, then pulsed with 1 mM H_2_O_2_ for 1 minute. The biotinylation signal was detected with fluorescein-conjugated streptavidin. The Cut6 and Pyr1 bands are in the same positions as in Fig.1 and tagged versions are indicated with an asterisk. Two identical samples for each strain were incubated with BP and activated with H_2_O_2_, and the identical samples were combined for immunodepletion and/or streptavidin purification as described in the text. *(**B**)* Coomassie blue staining of biotinylated proteins pulled down from samples with and without anti-Myc immunodepletion. Duplicate lysates of the *cut6^+^-13myc pyr1^+^-13myc* cells in panel A were first incubated with anti-Myc agarose beads, the combined supernatants were then incubated with streptavidin beads. The lysates of wild type *cut6^+^*and *pyr1^+^* non-depleted cells were directly incubated with streptavidin beads. The enriched proteins on the streptavidin beads were examined with Coomassie blue staining before subjected to TMT-mass spectrometry analysis. *(**C**)* Scatter plot showing the immunodepleted sample TMT intensities versus the non-depleted sample. The (x,y) coordinates of Cut6 and Pyr1 are (110.3,187.7) and (262.5,104.7), respectively. The data for this graph are in Supplemental Table 2.

To assess whether immunodepletion increased the identification of Pef1 neighbors, a ratio of the TMT tag signal for each of the identified proteins was calculated by dividing the signal of the immunodepleted BP-labeled sample by the non-depleted BP-labeled sample (Fig. 3C, Supplemental Table 2). The slope of the immunodepleted samples versus the non-depleted samples was negative, indicating that some low abundance peptides had stronger signals in the immunodepleted cells than in the control, non-depleted cells: 5.9% of the TMT identifiers by showed an increase of 2-fold or more in the immunodepleted sample, while 49.7% of the TMT identifiers decreased by at least 2-fold. Cut6 and Pyr1 were identified in both samples, with Cut6 being over-represented in the immunodepleted sample and Pyr1 being under-represented in the depleted sample (Fig. 3C), indicating that immunodepletion of the tagged proteins was not uniform. The reason for the negative slope for peptide abundance is not obvious and may reflect some non-specific protein loss during immunodepletion. As the number and identity of Pef1-APEX2-3V5 biotinylated proteins was not significantly changed by depleting Cut6 and Pyr1, subsequent experiments did not use immunodepletion.

### Identification of Pef1 neighbors in exponentially growing cells

To identify high-confidence Pef1 protein neighbors, two biological replicates with two types of control samples were used: *pef1^+^-APEX2-3V5* cells treated solely with BP and wild-type *pef1^+^* cells treated with both H_2_O_2_ and BP. Analysis of aliquots of the streptavidin-purified proteins by SDS-PAGE showed that more proteins were purified from the *pef1^+^-APEX2-3V5* sample treated with H_2_O_2_ (Fig. 4). The proteins on the streptavidin beads were then digested with trypsin, the released peptides were subjected to isobaric TMT-labeling and analyzed by mass spectrometry. A total of 264 *S. pombe* proteins were identified with 2 or more unique peptides (Supplemental Table 3, tab labeled Full Table). For each identified protein, the TMT ratio of the negative controls that lack biotinylation (Supplemental Table 3, tab labeled wt pef1 vs. pef1-APEX2 BP only) was 2.1 or less. Based on these criteria, specific proteins were not enriched in the absence of APEX2-based proximity labeling.

**Fig. 4.**
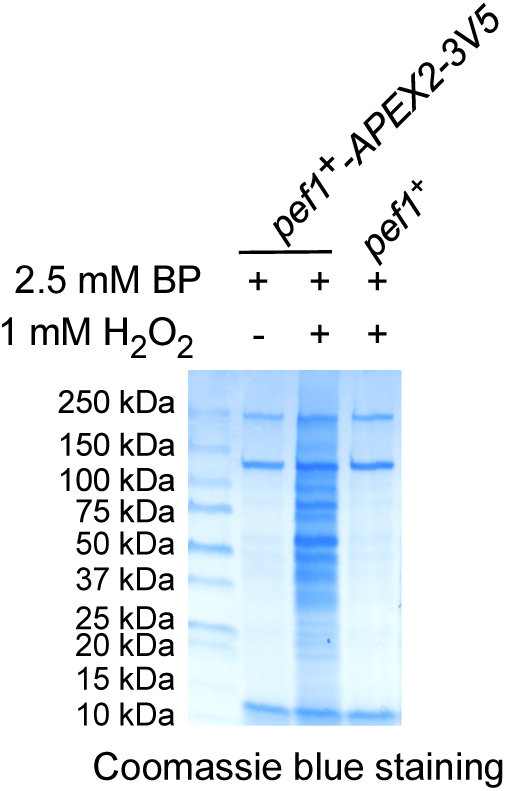
SDS-PAGE analysis of total biotinylated proteins of *pef1^+^-APEX2* cells and negative control cells lacking APEX2. The *pef1^+^-APEX2-3V5* cells treated with BP alone, or both BP and H_2_O_2_, and wild-type *pef1^+^* cells treated with both BP and H_2_O_2_ were incubated with streptavidin beads. Aliquots of streptavidin-bound proteins were examined using SDS-PAGE followed by Coomassie Blue staining. The image represents one biological replicate. Two biological repeats of each sample were subjected to TMT-mass spectrometry analysis.

A total of 264 proteins were identified in the *pef1^+^-APEX2-3V5* labeling. These proteins were filtered based on the following criteria: each protein must have at least two unique peptides identified, the TMT ratio of the experimental sample over each non-biotinylated control sample must be at least 2-fold, and the results should be consistent across two biological replicates. As a result, a total of 255 high-confidence neighbors of Pef1 were identified (Supplemental Table S3, tab labeled 255 IDs with a TMT ratio ≥2).

To elucidate the biological processes in which Pef1 neighbors participate, the 255 identifiers were categorized using Gene ontology biological process terms curated by PomBase (Supplemental Table 4) (31,32). The most frequent categories were cytoplasmic translation (70 genes), ribosome biogenesis (29 genes), and other cytoplasmic processes such as vesicle-mediated transport (19 genes) and actin cytoskeleton organization (9 genes). Nuclear processes such as chromatin organization, DNA-templated transcription and DNA repair were also identified (9, 12 and 2 genes, respectively). The proximity of Pef1 molecules near these proteins raises the possibility that Pef1 activity may regulate some these processes. Pef1 had not been previously associated with many of these GO terms, and the new link to DNA repair and DNA damage was examined.

### Pef1 physically interacts with Rad24 and modulates the response of *rad24Δ* cells to DNA damaging agents

The 14-3-3 protein Rad24 was consistently identified in Pef1-APEX2-3V5 labeled samples with multiple peptides (Supplemental Table 3, Full Table tab). Rad24 regulates the DNA damage response (39,40), while no role for Pef1 in this process was previously known, and physical or genetic interactions between Pef1 and Rad24 had not been previously identified (31). Rad24-Pef1 physical and phenotypic interactions were therefore tested.

To confirm the proximity relationship between Pef1 and Rad24, reciprocal proximity labeling was conducted. Rad24-APEX2-3V5 cells expressing either Pef1-3V5 or wild-type Pef1 were labeled with BP and H_2_O_2_, biotinylated proteins were purified using streptavidin beads, and the purified, biotinylated proteins were analyzed by Western blotting. The presence of the Pef1-3V5 band was clearly detectable in the proteins biotinylated by Rad24-APEX2 (Fig. 5A), indicating that Pef1 is a neighbor of Rad24.

**Fig. 5.**
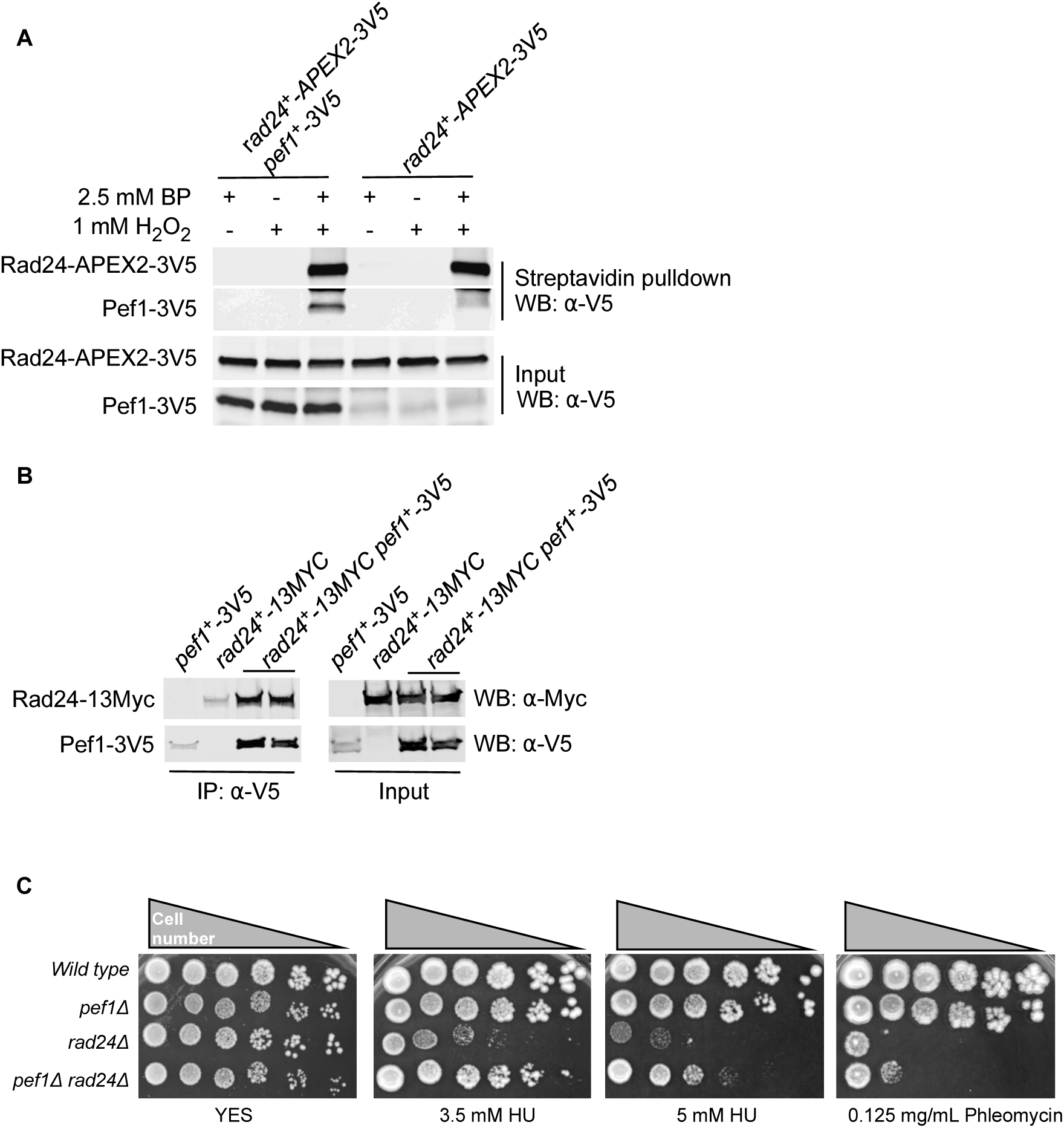
Proximity, physical and genetic interactions between Rad24 and Pef1. *(**A**)* Reciprocal proximity labeling. Genomic *rad24^+^* was tagged with APEX2, while genomic *pef1^+^* was either wild type or tagged with 3V5. The transcription of the both genes were driven by their endogenous promoters. Exponential cells were incubated with BP and Zymolyase for 30 minutes, pulsed with 1 mM H_2_O_2_ for 1 minute and biotinylated proteins were enriched with streptavidin beads and analyzed with Western blotting against V5. Either BP, or H_2_O_2_ were omitted in negative control samples. Input samples were aliquots (30 µg protein) of the extract that was subjected to incubation with streptavidin beads. *(**B**)* Rad24 co-immunoprecipitated with Pef1. Genomic *rad24^+^* was tagged with 13Myc; while genomic *pef1^+^* was tagged with triple V5. Lysate of exponential cells were incubated with anti-V5 agarose beads. The immunoprecipitated proteins were analyzed by Western blotting using antibodies against V5 or Myc. Duplicate precipitations from two *rad24^+^-13myc pef1^+^-3V5* extracts are shown. *(**C**)* Deletion of *pef1* partially rescued the sensitivity of *rad24Δ* cells to DNA damaging agents. Five-fold serial dilutions of exponentially growing cells were spotted onto YES plates with or without DNA damaging drugs and incubated at 32°C for 3 to 4 days and then imaged.

To test whether proximity labeling was a consequence of a physical interaction between Rad24 and Pef1, the proteins were tagged and tested in co-immunoprecipitation assays. The *rad24^+^* genomic locus was fused to the coding sequences for 13Myc epitopes while the *pef1^+^* locus was tagged with three V5 epitopes. Immunoprecipitates of Pef1-3V5 showed co-precipitation of Rad24-13Myc (Fig. 5B). Therefore, Rad24 and Pef1 physically interact.

To determine if the Pef1-Rad24 physical interaction may modulate the function of Rad24, the effect of eliminating Pef1 on Rad24 function were tested. Rad24 proteins play an important role in activating cell cycle checkpoints and initiating DNA damage repair (39,40), and cells lacking Rad24 are sensitive to DNA damaging agents such as hydroxyurea and phleomycin (41,42). The effect of eliminating Pef1 in cells that have or lack Rad24 on growth in presence of low levels of these DNA damaging agents was therefore tested. Serially diluted wild type, *rad24Δ*, *pef1Δ*, and *rad24Δ pef1Δ* cells were spotted onto YES plates with or without DNA damaging agents and incubated at 32°C. Wild type and *pef1Δ* cells grew the same on all media, and growth of the *rad24Δ* cells was inhibited by hydroxyurea and phleomycin. The *rad24Δ pef1Δ* double mutant grew more than the *rad24Δ* single mutant but less than the wild type and *pef1Δ* cells (Fig. 5C). These results indicate that Pef1 can modulate the cellular response to DNA damaging agents when Rad24 is absent.

### A known Pef1-interacting protein that is expressed at low levels was biotinylated by Pef1-APEX2

The proteins biotinylated by Pef1-APEX2 and identified by mass spectrometry did not include some of the proteins previously shown to physically interact with Pef1. We previously identified the135 kD threonine-serine kinase Cek1 as a Pef1-interacting partner (4), but no Cek1-derived peptides were identified in our experiments (Supplemental Table 3). Cek1 is expressed at very low levels in exponential cells (<200 molecules per cell (43,44)), so it was possible that the on-bead digestion and TMT-based multiplexing in our biotinylated samples was not sufficiently sensitive to detect this protein. To investigate whether Cek1 could be biotinylated by Pef1-APEX2-3V5, the *cek1^+^* ORF was tagged with a triple V5 epitope in pef1-APEX2-3V5 cells. After proximity labeling, the biotinylated proteins were purified on Streptavidin beads and analyzed by Western with an antibody against V5, and the Cek1 protein was detected (Fig. 6). Therefore, a known Pef1 interactor expressed at low levels was biotinylated by proximity labeling, but the current method and criteria for MS detection (TMT multiplexing and ≥ 2 peptides per protein) did not allow comprehensive detection of all Pef1 neighbors.

**Fig. 6.**
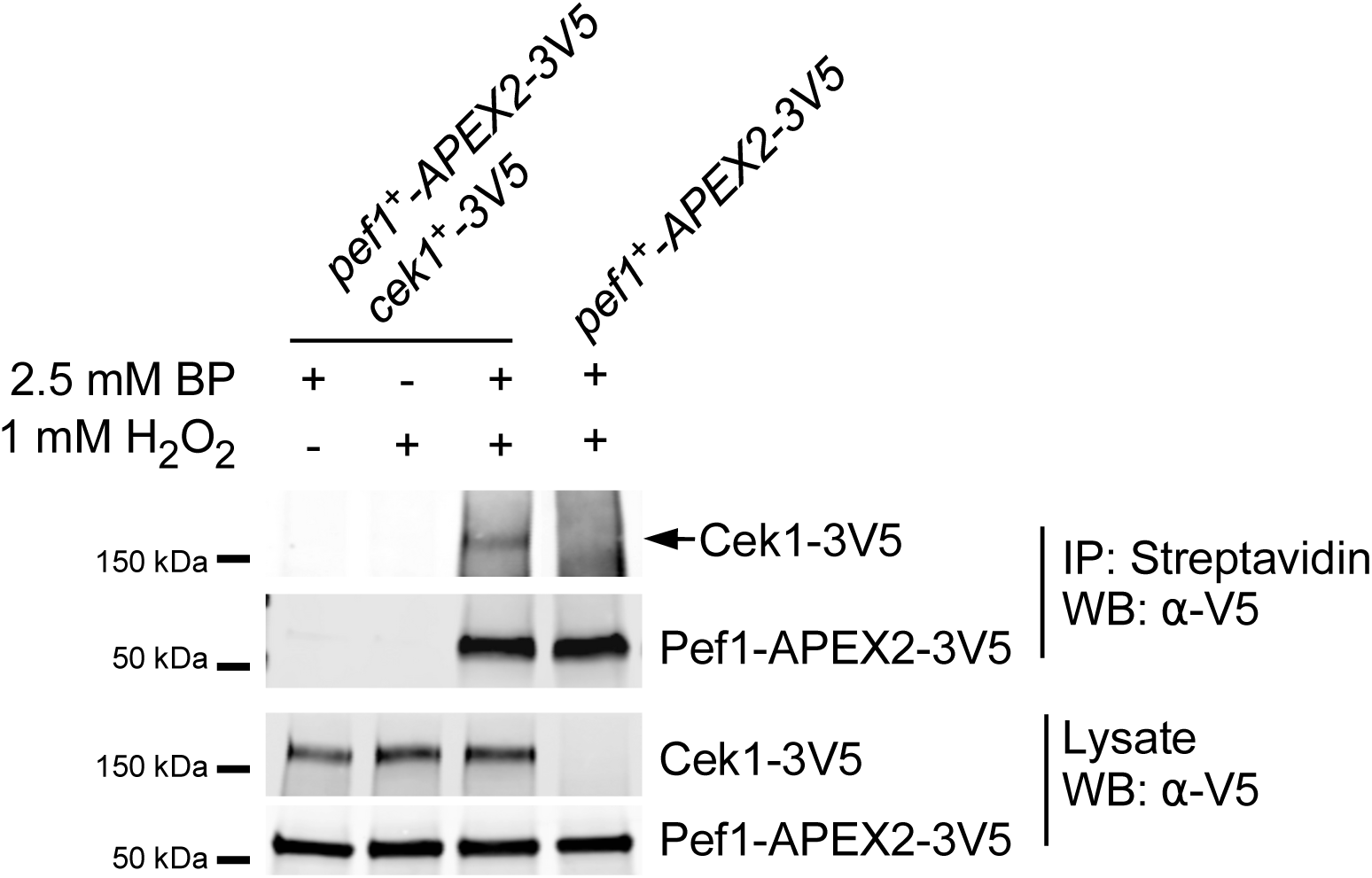
Labeling, purification and detection of the low abundance, Pef1-interacting protein Cek1 by Western blotting. Cells expressing *cek1^+^-3V5* and *pef1^+^-APEX2-3V5* or *pef1^+^-APEX2-3V5* alone from their genomic loci were treated with BP and H_2_O_2_ and lysates were affinity purified on streptavidin beads. Lysates and purified proteins were analyzed by analyzed by Western blotting using anti-V5 antibody. The streptavidin-purified Cek1-3V5 band is indicate by an arrow.

### Pef1-APEX2 proximity labeling in autophagic cells identifies 177 protein neighbors

The above results indicate that APEX2-based proximity labeling can identify novel Pef1-interacting partners in exponentially growing cells. As APEX2-mediated labeling can be rapidly induced, this approach was applied to cells induced for autophagy as the presence of Pef1 modulates autophagy levels ((8), our unpublished data). In fission yeast, autophagy can be induced by nitrogen starvation. In these experiments, exponential Pef1-APEX2-3V5 cells expressing the autophagy reporter GFP-Atg8 (9,45) were transferred to nitrogen-free medium and starved for 6 hours, followed by the proximity labeling protocol with or without the H_2_O_2_ pulse. Autophagy induction was confirmed by cleavage of GFP-Atg8 to produce free GFP in cell lysates and successful proximity labeling with BP in the H_2_O_2_ pulsed samples and was confirmed by Western blotting probed with streptavidin (Fig. 7).

**Fig. 7.**
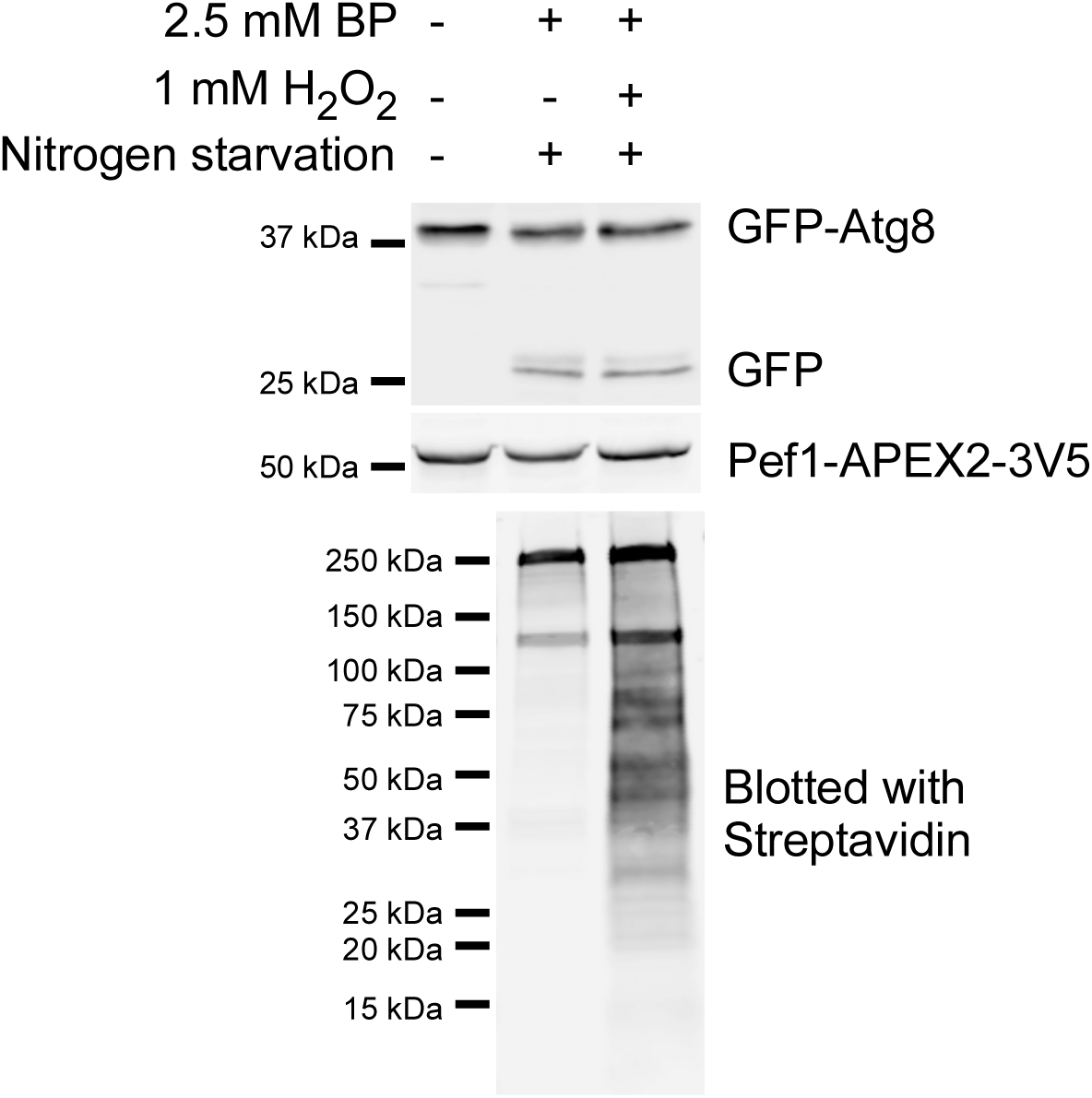
Proximity labeling in *pef1^+^-APEX2-3V5* cells induced for autophagy. Exponentially growing *pef1^+^-APEX2-3V5* cells harboring a GFP-atg8 expression plasmid were nitrogen starved for 6 hours, followed by proximity labeling. Autophagy induction was confirmed by detection of free GFP with anti-GFP antibody, using Pef1-APEX2-3V5 as a loading control by probing with anti-V5 antibody (upper panel). Proximity labeling efficiency was confirmed with Western blotting against fluorescein-conjugated streptavidin (lower panel).

Based on the results in exponential cells with TMT labeling and multiplexing of samples on mass spectrometry runs, a different approach to identify high-confidence protein neighbors was tested for autophagic cells. Pairs of proximity labeled and negative control samples (i.e. lacking the H_2_O_2_ pulse) were performed in triplicate, purified on streptavidin beads and each preparation was analyzed using mass spectrometry. Proteins identified with at least 2 unique peptides were considered genuine and retained for analysis. In the negative control samples, a total of 142 proteins, 236 proteins and 267 proteins were identified in the different experiments. Cut6 and Pyr1 were detected in these samples. The negative control samples therefore included the endogenously biotinylated proteins, and likely included proteins associated with the endogenously biotinylated proteins and proteins that non-specifically associate with the streptavidin beads. In the proximity labeled samples, a total of 240 proteins, 358 proteins and 366 proteins were identified in the different experiments (Supplementary Table 5). As in exponential cells, the low abundance Pef1 physically interacting protein Cek1 was not detected by mass spectrometry. Label-free quantitation (LFQ) intensities were utilized to indicate the relative amounts of the peptides and to represent the ratio changes of each protein across the proximity labeled and control samples. All proteins were filtered using the criteria that the LFQ ratio of the proximity labeled sample over its control sample was no less than 1.5 and was repeated in at least two biological repeats. Application of these criteria resulted in a total of 177 high confidence Pef1 neighbors in autophagic cells (Supplementary Table 5). This filtering process eliminated the Cut6 and Pyr1, as these endogenously biotinylated proteins were present in the positive and negative controls at similar levels.

The 177 Pef1 neighbors in in autophagic cells were classified using GO biological process terms (Supplemental Table 5, tab labeled Bio. Proc. GO slim). The top three over-represented biological processes in autophagic cells are cellular amino acid metabolic process (enriched with 17% identifiers, 30 out of 177), nucleobase-containing small molecule metabolic process (enriched with 15% identifiers, 27 out of 177) and cytoplasmic translation (enriched with 12% identifiers, 22 out of 177). The top 25 GO terms for both autophagic and exponentially growing cells were very similar, with small differences in percentages of the total number of proteins for some categories which only had 1 to 3 proteins identified (Supplemental Table 6, tab labeled GO Slim comparison).

While the identified GO terms were very similar for the exponential and autophagic cell-identified proteins, each experiment also identified specific proteins. While 96 proteins were identified in both experiments, the exponential cells identified 159 proteins not seen in autophagic cells experiments (Supplemental Table 6, exponential_specific tab). The largest classes of unique proteins in exponential cells involved cytoplasmic translation, ribosomal subunits and amino acid metabolism, suggesting a role for Pef1 in modulating protein production. The proteins identified in exponential cells also included Cut6 and Pyr1, which were filtered out of the autophagic protein list.

The autophagic cells identified 81 Pef1 neighbors not seen in the exponential cell experiments (Supplemental Table 6, autophagic_specific tab). Of the unique autophagic cell proteins, 2 proteins were associated with autophagy GO term: Vtc4, a vacuolar transporter chaperone and polyphosphate synthase subunit, and Osh2, a sterol transfer protein. Three other proteins with the autophagy GO term that were present in both exponential and autophagic cells were Cdc48 (an ATPase involved in ubiquitin-mediated degradation), Scs2 (a protein related to autophagy of the ER) and Osh6 (another sterol transfer protein). In addition to these autophagy-related proteins, proteins related to vesicle-mediated transport, a process that overlaps with transporting proteins to the vacuole, were identified in the exponential cells, autophagic cells and both samples (8, 11 and 11 proteins respectively, Supplemental Table 6). Consequently, APEX2-mediated proximity labeling was able to identify potential targets by which Pef1 may regulate autophagy.

## DISCUSSION

To reveal the regulatory processes directly impacted by the non-canonical cdk Pef1, an ortholog of human cdk5 and *S. cerevisiae* Pho85, we optimized APEX2-mediated proximity labeling in the fission yeast *S. pombe*. A major issue was to determine which conditions allowed APEX2-mediated biotinylation of multiple proteins in different subcellular compartments. Conditions that allow normal growth, such as presence of glucose or reduced nitrogen in the medium, inhibited biotinylation, possibly be preventing biotin-phenol uptake by the cells. This study used 30 minutes in buffered 1.2 M sorbitol with a cell wall digesting enzyme (Zymolyase) in the presence of biotin-phenol, which gave the highest level of biotinylation for all conditions tested (Figs. 1, 2).

It is worth noting that conditions with yeast synthetic medium (EMM) lacking glucose or reduced nitrogen in 1.2 M sorbitol and an increased amount of Zymolyase also resulted in substantial labeling, indicating that this modified EMM-1.2 M sorbitol medium may be useful in some experiments. A second aspect was testing different mass spectrometry approaches to identify Pef1 protein neighbors. Proximity labeling generates large lists of candidate proteins that may interact with the protein fused to APEX2, and this work compared TMT-multiplexed and single protein mass spectrometry approaches for generating these lists from biotinylated proteins digested on streptavidin beads. Both approaches generated large lists of proteins, indicating that the method of choice depends upon the availability and cost of mass spectrometry time. An outcome of the Pef1-APEX2 model system was the identification of a novel Pef1 interaction with Rad24 and its paralog Rad25, interactions which were not predicted by the existing literature.

Reciprocal proximity labeling, co-immunoprecipitation and genetic experiments established a role for Pef1 in affecting cell growth in response to DNA damaging agents when Rad24 was absent (Fig. 5). How Pef1 and Rad24 modulate the function of each protein and cell growth is currently unknown.

Validation that Rad24 was a Pef1 neighbor was first done by reciprocal proximity labeling (i.e. Rad24-APEX2 biotinylation of Pef1) for several reasons. The Pef1 kinase may only transiently interact with its substrates, which would allow proximity labeling but might not be stable enough for co-immunoprecipitation. Another possibility is that if Pef1 interacts with one protein in a larger complex, and the requirement for unmodified Lys, Cys or Tyr residues for APEX2 labeling (46) may limit which proteins in a complex are labeled, but interaction between complex members may not be strong enough to support co-immunoprecipitation. Consequently, while co-immunoprecipitation is strong evidence for physical interaction, a negative result would not mean that the APEX2-fused protein does not regulate or associate with the identified neighbor protein. Since co-immunoprecipitation in yeast generally requires making an epitope-tagged protein, creating the APEX2 fusion can be done at the same time. Therefore, reciprocal proximity labeling experiments are important to validate if the APEX2-fused protein and the biotinylated protein are in close proximity.

Pef1-APEX2 labeling identified many new Pef1 neighbors, but labeling was not comprehensive as known interacting proteins such as the Pef1 cyclins Pas1, Clg1 or Psl1 and the associated downstream kinase Cek1 were not identified (4,47). All of these proteins are estimated to be present at <3000 copies/cell whereas identified proteins such as Rad24, Cdc48 or Osh6 (Supplementary Table S6, Common_Autophagy-Exponential tab) are present at >15,000 copies/cell (31). We found that Cek1 was biotinylated in Pef1-APEX2 cells by Western analysis of streptavidin-purified proteins (Fig. 6), suggesting that detection by mass spectrometry required a threshold protein level that Cek1 did not meet. Protein digestion was performed on streptavidin beads to obtain identification with the shortest amount of mass spectrometry time. An alternative approach that may be more sensitive would be to release the biotinylated proteins from the beads by boiling and analyzing the eluant by SDS-PAGE. Cutting the gel lane with the eluant into pieces and digesting and analyzing peptides present from each piece would reduce the total number of peptides and may increase the ability to detect low abundance proteins. However, given the number of experiments and controls to be analyzed, the cost and practicality of this approach may be limited.

Two approaches for mass spectrometry analysis of the multiple experimental and control samples for each experiment were used to compare the number of identified proteins. For exponential cells, the proximity labeled and negative control samples were digested on beads, and normalized amounts of peptides were coupled to TMT isobaric tags so several experiments could be analyzed in one multiplexed run and the source of the detected peptides identified by the attached TMT tag. The peptides from the autophagic cell triplicate experiments and triplicate negative controls were each analyzed in a single mass spectrometry run and compared directly. Both approaches identified similar numbers of Pef1 neighbors before filtering for high confidence protein identification. Neither approach detected Cek1, suggesting no major advantage to either method for detecting low abundance proteins. The difference in the final number of proteins identified in the exponential and autophagic cell experiments was due to the filtering method. In autophagic cells, proteins identified in the negative controls were excluded from the list of Pef1-APEX2 labeled cells, which eliminated the endogenously biotinylated Cut6 and Pyr1 proteins and may have removed proteins that bind non-specifically to the streptavidin beads. In contrast, the approach in exponential cells used the relative amounts of TMT peptides to determine whether proteins were identified as Pef1 neighbors. Both approaches identified the novel Pef1-interacting protein Rad24 (Supplementary Table S6, Common_Autophagy-Exponential tab), but the TMT filtering approach may not have been sufficiently strict. We note that the ratio of TMT abundance for Rad24 peptides in the labeled samples over the different unlabeled controls was ∼50 or 78, while the ratio for the endogenously biotinylated proteins Cut6 and Pyr1 were both less than 4 (Supplementary Table S4, Full Table tab). The ratio for Cut6 and Pyr1 peptides in a TMT-based approach may therefore serve as the lowest cutoff for high confidence protein neighbors. This cutoff would make the number of protein neighbors identified in the two approaches much closer, with the TMT multiplexing approach reducing the amount of required mass spectrometry time.

The Pef1 neighbors identified in exponential and autophagic cells strongly suggest that *S. pombe* Pef1 regulates actin cytoskeleton dynamics, similar to the evolutionarily distant human cdk5 and *S. cerevisiae* Pho85 (48). Actin (Act1), the actin filament depolymerization factor Cofilin (Adf1), and Pan1 and Ede1 of the complex that links actin to endocytic vesicles were identified as a Pef1 neighbors in both exponential and autophagic cells (Supplementary Table S7). Arp2 and Arc2, part of a complex that allows for actin filament branching (49–52), were identified in autophagic cells. Pef1 regulates autophagy, and actin filament dynamics are important for various stages of autophagy (53–56), and these neighbors may reveal how Pef1 regulates this process.

Autophagy is a membrane trafficking process that involves assembly of the autophagosomes from phagophore assembly sites in the ER (called omegasomes in the mammalian cells) and eventual fusion of the double-membraned autophagosome with the yeast vacuole (the lysosome in mammalian cells) (9,57,58), and multiple Pef1 neighbors have roles in vesicular trafficking.

The COPII vesicle components Sec31 and Sec21 were identified in both exponential and autophagic cells, and components of the secretory machinery, Sec24 and Ypt1 in exponential cells and Sec18 and Ypt3 in autophagic cells were also identified (Supplemental Table 7). A vesicle-associated protein involved in ER membrane recycling (reticulophagy), Scs2, was identified in both exponential and autophagic cells, consistent with a link to autophagy.

Trafficking of the autophagy protein Atg9 relies on clathrin-dependent endocytosis (59,60), and Pef1 neighbors involved in endocytosis in exponential and/or autophagic cells include clathrin heavy chain Chc1, clathrin interactors Ent1, Ent3, End4, endocytosis-related proteins Pan1 and Ede1 and the endocytic membrane scission protein Hob1 (Supplemental Table S7), all of which are conserved in mammals (61). Some of these *S. pombe* associations are likely functional, as a human homolog of Hob1, amphilysin I, is a substrate of cdk5 (62–64). These results imply that Pef1 can modulate vesicular trafficking during normal cell growth and may regulate it during autophagy.

The proximity labeling approach that identified a new role for Pef1 in DNA repair and suggests a role in vesicular trafficking used the originally described biotin-phenol as the labeling substrate (11). This approach in *S. pombe* required cell wall digestion, osmotic support and excluding nitrogen and glucose for optimal labeling (Fig. 2). Alternative methods and substrates for APEX2 have been recently developed that may alter these requirements. One alternative used in *S. cerevisiae* was to semi-permeabilize cells by freezing them in a 15% glycerol medium at -80°C, thawing the cells and adding biotin-phenol, which allowed APEX2-mediated biotinylation (65). New APEX2 substrates that label nucleic acids may also be useful for *S. pombe.* An alkyne-phenol alternative substrate used in *S. cerevisiae* with an intact cell wall was able to enter cells better than biotin-phenol and allowed subsequent biotinylation using click-chemistry in cell lysates (66). This substrate may reduce the amount of cell wall digestion needed in *S pombe* to achieve labeling. Biotin-aniline and biotin-naphthylamine have shown promise for labeling RNA and mapping the mitochondrial transcriptome (67). The hydrophobic nature of these substrates may facilitate crossing the yeast cell wall, similar the alkyne-phenol substrates. These alternative approaches have high potential for probing protein and RNA neighbors of a protein of interest, even in the more challenging situation of organisms with a cell wall. Optimizing these approaches will likely require a systematic analysis of conditions that allow maximum labeling, as was done for biotin-phenol (Fig. 2). The successful application of APEX2-mediated labeling to *S. pombe* in this work and advances in substrate design will greatly facilitate experiments for discovering new functions for evolutionarily-conserved proteins in this model organism that will impact our understanding of similar processes in mammals and other species.

## DATA AVAILABILITY

The authors confirm that the data supporting the findings of this study are available within the article or its supplementary materials.

## SUPPLEMENTAL MATERIAL

Supplementary material are 5 separate tables in Excel format.

## CONTRIBUTIONS

Experimental design: H. Zhang, B. Willard and K. Runge. Performed the biological experiments: H. Zhang. Performed the MS sample preparation and analysis: D. Zhang, L. Li and B. Willard. Analyzed the data: H. Zhang, D. Zhang, B. Willard, K. Runge. Wrote the paper: H. Zhang, K. Runge.

## ACKNOWLEDGEMENTS

We thank Drs. Kathleen L. Berkner and Julien Audry for helpful discussions during the course of this work and Carly Kerr for technical support.

---

**Supplemental Table 1.** Yeast strains used in this work.

| Strain number | Genotype | Source | Fig. |
| --- | --- | --- | --- |
| KRP1 | *h<sup>-</sup>, ade6-M216 ura4-D18 leu1-32 his7-366* | Lab stock | Fig. 1 and 3 |
| KRP2 | *h<sup>+</sup>, ade6-M216 ura4-D18 leu1-32 his7-366* | Lab stock | Fig. 3 |
| KRP131 | *h<sup>-</sup> ade6-M216 ura4-D18 his7-366 pef1Δ::ura4<sup>+</sup>* | Lab stock (4) | Fig. 5 |
| KRP617 | *h<sup>-</sup>, ade6-M216 ura4-D18 leu1-32 his7-366 pef1::pef1-APEX2-3V5-KanMX* | This study | Fig. 1, 2, 3, 4, 6 |
| KRP622 | *h<sup>-</sup>, ade6-M216 ura4-D18 leu1-32 his7-366 pef1::pef1-APEX2-3V5-KanMX*<br>*pyr1::pyr1-13myc-ura4* | This study |  |
| KRP623 | *h<sup>+</sup>, ade6-M216 ura4-D18 leu1-32 his7-366 cut6-13myc-ura4* | This study |  |
| KRP618 | *h<sup>-</sup>, ade6-M216 ura4-D18 leu1-32 his7-366 pef1::pef1-APEX2-3V5-KanMX*<br>*pyr1::pyr1-13myc-ura4 cut6::cut6-13myc-ura4* | This study | Fig. 3 |
| KRP623 | *h<sup>-</sup> ade6-M216 his7-366 rad24Δ::KanMX* | This study | Fig. 5 |
| KRP624 | *h<sup>-</sup> ade6-M216 ura4-D18 his7-366 pef1Δ::ura4<sup>+</sup> rad24Δ::KanMX* | This study | Fig. 5 |
| KRP629 | *h<sup>-</sup>, ade6-M216 ura4-D18 leu1-32 his7-366 pyr1::pyr1-13myc-ura4 cut6::cut6-13myc-ura4*<br>*rad24::rad24-APEX2-3V5-natMX* | This study | Fig. 5 |
| KRP630 | *h<sup>-</sup>, ade6-M216 ura4-D18 leu1-32 his7-366 pyr1::pyr1-13myc-ura4 cut6::cut6-13myc-ura4*<br>*rad24::rad24-APEX2-3V5-natMX pef1-3V5-KanMX* | This study | Fig. 5 |
| KRP631 | *h<sup>-</sup>, ade6-M216 ura4-D18 leu1-32 his7-366 pef1::pef1-3V5-natMX* | This study | Fig. 5 |
| KRP633 | *h<sup>-</sup>, ade6-M216 ura4-D18 leu1-32 his7-366 pef1::pef1-3V5-natMX rad24::rad24-13myc-kanMX* | This study | Fig. 5 |
| KRP634 | *h<sup>-</sup>, ade6-M216 ura4-D18 leu1-32 his7-366 rad24::rad24-13myc-kanMX* | This study | Fig. 5 |
| KRP635 | *h<sup>-</sup>, ade6-M216 ura4-D18 leu1-32 his7-366 pef1::pef1-APEX2-3V5-kanMX*<br>*cek1::cek1-3V5-natMX* | This study | Fig. 6 |
| KRP642 | *h<sup>-</sup>, ade6-M216 ura4-D18 leu1-32 his7-366 pef1::pef1-APEX2-3V5-KanMX*<br>*pREP81-GFP-atg8* | This study | Fig. 7 |

**Supplemental Table 7.** Pef1 neighbors related to actin and vesicular trafficking. Some proteins function in both processes.

| **experiment** | **process** | **actin**<br>**cytoskeleton** | **vesicular trafficking** |
| --- | --- | --- | --- |
| exponential and autophagic cells |  | Act1, Adf1, Pan1, Ede1 | Sec31, Sec21, Chc1, Ede1, Ent3, Hob1, Pan1, Scs2 |
| exponential cells only |  | Shd1 | Sec24, Ypt1 |
| autophagic cells only |  | Arp2, Arc2 | Sec18, Ypt3, Ent1, End4 |

